# Mapping Tree Cover Expansion in Montana, U.S.A. Rangelands Using High-Resolution Historical Aerial Imagery

**DOI:** 10.1101/2022.12.28.522142

**Authors:** Scott Morford, Brady W. Allred, Eric R. Jensen, Jeremy D. Maestas, Kristopher R. Mueller, Catherine L. Pacholski, Joseph T. Smith, Jason D. Tack, Kyle N. Tackett, David E. Naugle

## Abstract

Worldwide, trees are colonizing rangelands with high conservation value. The introduction of trees into grasslands and shrublands causes large-scale changes in ecosystem structure and function, which have cascading impacts to ecosystem services, biodiversity, and agricultural economies. Satellites are increasingly being used to track tree cover at continental to global scales, but these methods can only provide reliable estimates of change over recent decades. Given the slow pace of tree cover expansion, remote sensing techniques that can extend this historical record provide critical insights into the magnitude of environmental change. Here, we estimate conifer expansion in rangelands of the northern Great Plains, United States, North America, using historical aerial imagery from the mid-20th century and modern aerial imagery. We analyzed 19.3 million hectares of rangelands in Montana, USA, using a convolutional neural network (U-Net architecture) and cloud computing to detect tree features and tree cover change. Our bias-corrected results estimate 3.0 ± 0.2 million hectares of conifer tree cover expansion in Montana rangelands, which accounts for 15.4% of the total study area. Overall accuracy was greater than 91%, but producer accuracy was lower than user accuracy (0.60 vs. 0.88). Nonetheless, the omission errors were not spatially correlated, indicating that the method is reliable for estimating the spatial extent of tree cover expansion. Using the model results in conjunction with historical and modern imagery allows for effective communication of the scale of tree expansion while overcoming the recency effect caused by shifting environmental baselines.

## 1.0 Introduction

Tree cover expansion (hereafter, tree expansion) is a widespread global change phenomenon that alters rangeland ecosystems’ structure, function, and biodiversity (Nackley et al., 2017; Van Auken, 2009). Tracking local and regional tree expansion has been an active area of remote sensing research for more than 20 years, but only recently have operational products been developed to track tree cover change in rangelands at continental to global scales (Allred et al., 2021; Asner et al., 2003; Brown et al., 2022). Renewed concern regarding the accelerating degradation of global rangelands and loss of rangeland-obligate fauna highlight the need for tools and approaches to better track and communicate the extent to which tree expansion is modifying grassland and shrubland ecosystems (Bardgett et al., 2021; Lees et al., 2022).

Addressing tree and shrub encroachment in rangelands (defined as grasslands, shrublands, savannas, and open woodlands) is a globally important strategy for climate adaptation and biodiversity protection (Smith et al., 2022). In temperate zones of the northern hemisphere, managing grassland tree expansion also helps mitigate climate change by maintaining or restoring high land-surface albedo (Ge & Zou, 2013; Mykleby et al., 2017; Nuñez et al., 2021). Yet, the broader global change community shows sparse awareness of how increasing tree cover impacts the function and biodiversity of grasslands. This awareness gap is perhaps best illustrated by recent calls for afforestation of global grasslands to enhance land carbon sequestration (Bastin et al., 2019; Veldman et al., 2019).

This awareness gap extends to the wider public. Tree encroachment unfolds slowly over decades, and humans have difficulty perceiving incremental environmental change (Essl et al., 2015). Further, innate or cultural aesthetic preferences for trees in the environment coupled with high-profile campaigns promoting tree planting as a climate change solution also hinder the recognition of tree expansion and its potential to negatively impact the function of rangeland ecosystems (Cook & Cable, 1995; Han, 2007).

Providing scientists, land managers, and the public with a tool to visualize tree expansion is a powerful conservation communication strategy to overcome the recency effect associated with shifting environmental baselines (Jones et al., 2020; Soga & Gaston, 2018). While time-series analysis of satellite data provide robust spatiotemporal estimates of tree expansion, these moderate-resolution data products do little to visually convey how tree expansion is reorganizing the structure of grasslands as they are converted to woodlands and forests. In contrast, using historical aerial imagery allows users to see how tree cover expansion has unfolded at resolutions in-line with modern aerial and satellite mapping technologies.

This work builds upon recent innovations in photogrammetry, deep learning, and cloud-based geospatial processing to analyze tree cover change at previously impossible spatial and temporal scales. New photogrammetry techniques that utilize Structure from Motion (SfM) and MultiView Stereo (MVS) algorithms can efficiently georectify thousands of historical images with relatively little user intervention (Hirschmuller, 2008; Toldo et al., 2015). Deep learning approaches, such as convolutional neural networks, are now widely used across disciplines for automated feature detection and image segmentation (Ma et al., 2019). For example, the U-net architecture and its permutations are particularly effective at pixel-wise segmentation and are widely used in biomedical and environmental applications (Ronneberger et al., 2015; Wagner et al., 2019). Fusing and processing these data in cloud geospatial platforms, such as Google Earth Engine, provide efficient and scalable computing power that allow for these data to be analyzed by scientists without extensive expertise in high-performance computing.

Recent tree expansion work from the United States (U.S.) shows that tree cover in grasslands has increased by 85% over the past 30 years, resulting in 147,700 km^2^ of tree-free grasslands being converted to woodlands (Morford et al., 2022a). These changes to ecosystem structure rival the impacts from cropland conversion and drive significant losses of habitat for sagebrush- and grassland-obligate birds such as greater sage-grouse (*Centrocercus urophasianus*) and lesser prairie chickens (*Tympanuchus pallidicinctus*) (Baruch-Mordo et al., 2013; Lautenbach et al., 2017). The broader impacts of tree expansion to ecosystem services and biodiversity have been widely documented across the southern U.S. Great Plains where tree expansion has expanded rapidly over the past century. However, tree expansion has received relatively little attention in the northern Great Plains (Symstad & Leis, 2017). Current satellite observations and dynamic vegetation modeling report that tree expansion is rapidly increasing across the northern Great Plains and will continue to accelerate the conversion of grassland to woodlands in the coming decades as a result of climate change (Klemm et al., 2020; Shafer et al., 2015).

Here we analyze tree expansion over a 70+ year period in 19.2 million hectares of rangelands in the state of Montana, United States of America. We use high-resolution (< 2 meter ground sampling distance) historical and modern imagery to quantify the aerial extent of tree expansion since the mid-20th century. We also provide complimentary estimates for rangeland tree cover losses to help understand the magnitude and direction of ecosystem evolution between grasslands, woodlands, and forests.

We focus on Montana U.S.A. rangelands because they are a focal area for ongoing multinational conservation efforts to protect North American grasslands and their biodiversity (Epstein et al., 2021). Grasslands in this region are critical habitat for collapsing North American grassland bird populations, many species of which are highly sensitive to even low abundances of trees (Rosenberg et al., 2019). Additionally, tree encroachment in this region can lead to losses in forage production, which can exacerbate economic stress on small ranching operations and further promote land use conversion to row-crop agriculture, residential sub-division, and fossil-fuel energy development (Allred et al., 2015; Lark, 2020; Morford et al., 2022a).

## 2.0 Methods

We used high-resolution historical aerial imagery sourced from the United States Geological Survey (USGS) single-frame archive, modern aerial imagery from the National Agricultural Inventory Program (NAIP) and auxiliary land cover data to analyze tree expansion in Montana rangelands. Conceptually, we use pixel-wise segmentation in the historical and modern imagery to identify tree features in the imagery and then perform kernel smoothing to approximate tree density at scales of 0.40 hectares (one acre). We then calculate the difference between the two cover density products to identify areas of tree expansion or tree cover loss (**Fig. 1**).

**Figure 1:**
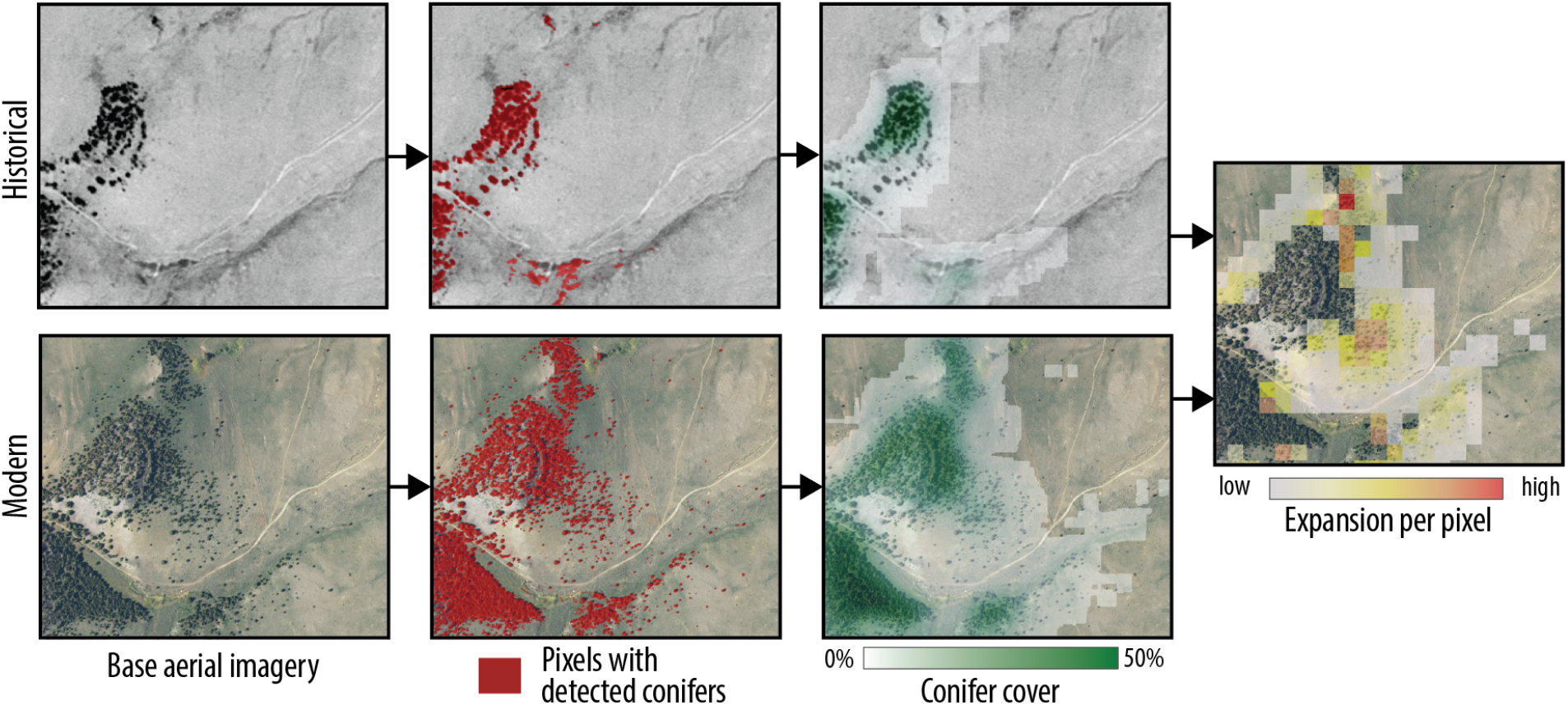
Conceptual workflow for identifying areas of tree encroachment using historical and modern aerial imagery.

We acquired high-resolution aerial images from the USGS archive collected by the U.S. Army between 1946 and 1977. The median year for imagery acquisition was 1954. The images were scanned by the USGS at a resolution of 25 micron (1,000 dots per inch) in 8-bit grayscale. Depending on acquisition altitude, these images produced digital orthoimagery with a ground sampling distance (GSD) between 0.8 and 2.0 meters (m). We exported the imagery at a GSD of 1.0 m for tree expansion analysis. The modern aerial imagery was sourced from 2019 and 2020 and was provided by NAIP as 4-band (RGB-IR) 8-bit images with a GSD of 0.6 m.

**Figure 2** presents the data-processing pipeline. The pipeline is segmented into three distinct workflows including orthorectification, feature detection, and spatial analysis.

**Figure 2.**
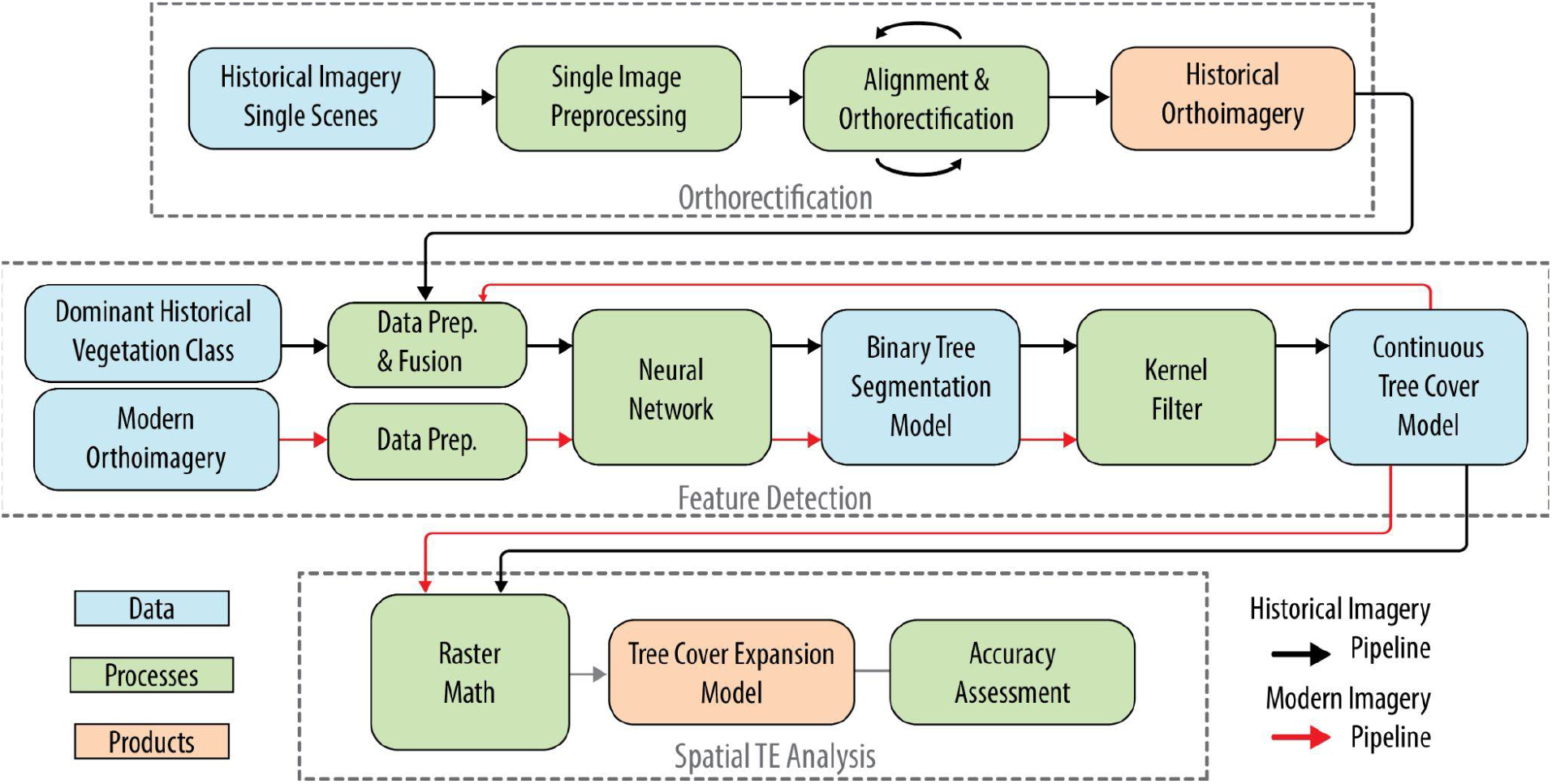
Data processing workflow. Our approach comprises of three modules: orthorectification of historical imagery, feature detection and mapping using U-net neural network, and spatial analysis of tree cover expansion and loss.

### 2.1 Orthorectification

We used MATLAB, Metashape, and QGIS to produce the historical orthoimage. Images from individual flight projects as identified by the USGS were processed together and then projects were merged together into a final seamless orthoimagery product. In MATLAB, we cropped the images to remove the film borders and applied a dehazing and low light filter to reduce vignetting and enhance image contrast. In Metashape, preliminary bundle adjustment was performed using image coordinates provided by USGS metadata and internal and external calibration parameters were determined by the software. Orthorectification used a DEM derived from MVS dense-point-cloud processing of the imagery.

After generating a preliminary orthoimage, we imported the imagery into a GIS and identified common features in historical and modern aerial imagery for use as ground control points (GCP). The GCPs and coordinates were then added to Metashape and the project was realigned using GCP coordinates rather than coordinates from the USGS metadata. Candidate orthoimagery products were then re-evaluated in a GIS to assess map alignment. If needed, additional GCP were added and the imagery was reprocessed until the horizontal offset between the historical and modern aerial imagery was on the order of 5 - 10 m.

### 2.2 Feature Detection

We applied the U-Net architecture as described in Ronneberger et al. (2015) to detect tree pixels in the aerial imagery. At a high level, the architecture includes a set of encoder blocks to detect tree features and decoder blocks to provide pixel-wise predictions for tree features. The architecture also uses skip connections to transmit spatial features from the encoder blocks to decoder blocks that may otherwise be lost during image downsampling (**Fig. 3**).

**Figure 3:**
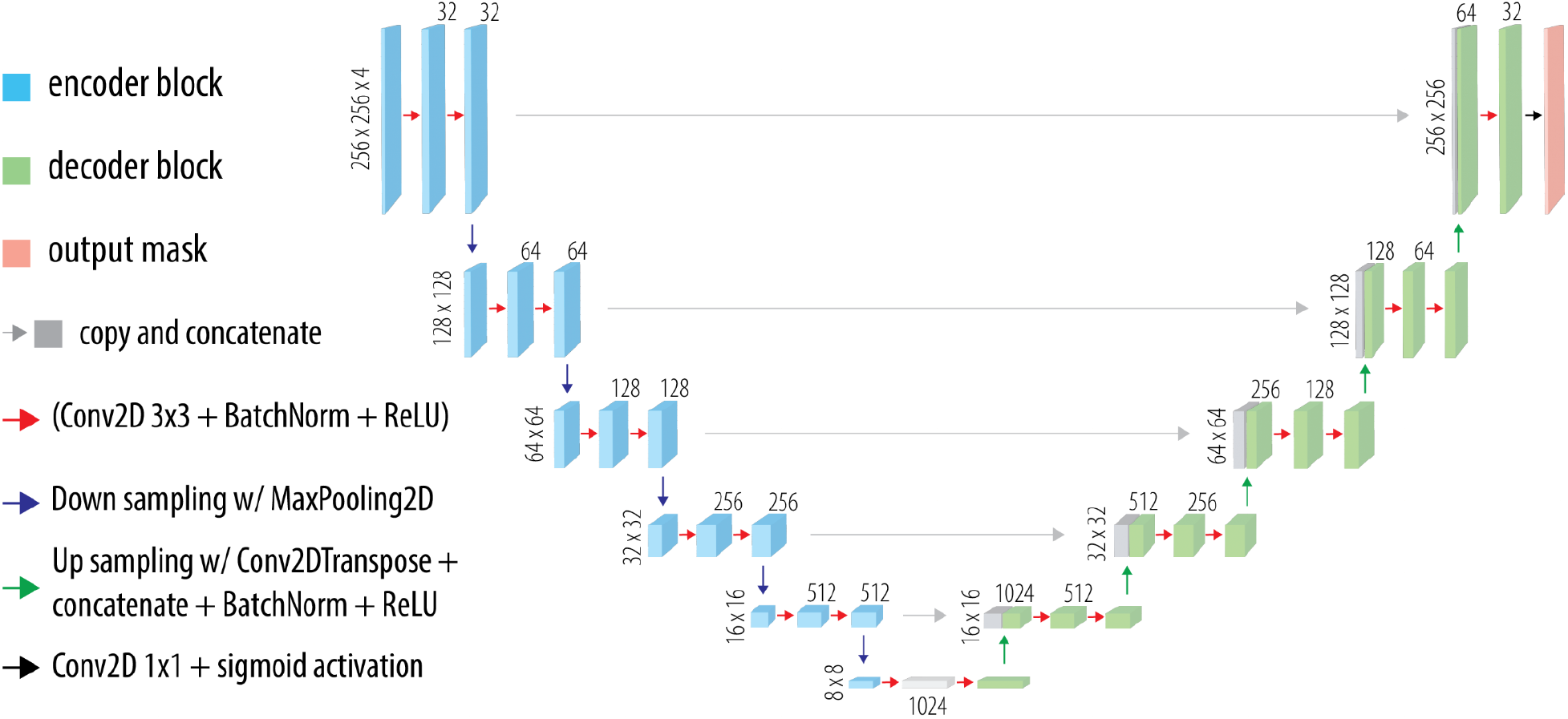
U-Net architecture used for tree feature detection in historical imagery.

For this analysis, the input channels, array size, and hyperparameter selection differed between the historical and modern imagery models due to differences in imagery inputs, processing constraints, and training performance (**Table 1**). Figure 3 presents the U-Net architecture used for historical imagery with an input image array of size 256 x 256 x 4.

**Table 1:** Tree Feature Detection model and Performance Metrics

|  | Historical Imagery | Modern Imagery |
| --- | --- | --- |
| **Tree feature detection** |  |  |
| Architecture | U-net | U-net |
| Input Image Dimensions | 256 X 256 X 4 | 1024 X 1024 X 4 |
| Channel inputs | - Grayscale Image | - Red channel |
|  | - GLCM Variance | - Green channel |
|  | - BPS Vegetation Class | - Blue channel |
|  | - Modern Tree Cover | - NDVI |
| Ground sampling distance | 1.0 meters | 0.6 meters |
| Total Model Parameters | 31,126,785 | 31,138,305 |
| Training images | 8,245 | 10,690 |
| Training pixels | 540 million | 11.2 billion |
| Optimizer | Adam | RMSProp |
| Loss | Dice coefficient | Dice coefficient |
| **Performance Metrics from Validation Dataset** |  |  |
| Dice | 0.9018 | 0.8141 |
| Jacard (IoU) | 0.8669 | 0.7311 |
| Sensitivity | 0.9858 | 0.9314 |
| Specificity | 0.9855 | 0.9769 |

The output was a 256 x 256 x 1 array where pixel values ranged from 0 to 1 indicating the probability of absence or presence of trees, respectively. The modern imagery model architecture takes the same form, but contains more encoder/decoder blocks due to its larger input size (1024 x 1024 x 4) necessitating additional downsampling and upsampling steps.

#### 2.2.1 Training data development

Utilizing GIS and Google Earth Engine, training data were created by manually annotating historical and modern imagery. Field data were not used. Pixels were visually evaluated at 1:1000 to 1:5000 scale to classify conifer species as the model target. We attempted to constrain our tree model detection to conifer trees because tree expansion in the northern Great Plains is driven by three conifer species including Douglas Fir (*Pseudotsuga menziesii*), Eastern Red Cedar (*Juniperus virginiana*) and Rocky Mountain Juniper (*Juniperus scopularum*). In modern aerial imagery, visually distinguishing between conifer and deciduous trees was relatively easy owing to differences in texture and color. Marking the historical imagery was more difficult because distinguishing between conifer and deciduous trees relied primarily on detecting textural differences in imagery and reliance on auxiliary data such as landscape position and vegetation classification.

We limited our training data to upland grassland and forested locations and omitted riparian, wetland, cropland, and urban areas using modern land cover classification from the USDA NASS, USGS NLCD, and Montana Natural Heritage Land Cover product. Locations for training data were stratified roughly equally among primary forests, woodlands, and rangeland cover classes, with an emphasis on capturing the transition between these cover types. For modern imagery, we labeled a total of 10,690 (1024 x 1024) tiles; for historical imagery we labeled 8,245 tiles (256 x 256). We used a 80% - 20% training validation split during model fitting.

#### 2.2.2 Data preparation and fusion

Inputs for the modern (2019) tree detection model were derived entirely from the 4-band NAIP imagery. We replaced the near-infrared channel (band 4) with NDVI calculated from the red and near-infrared bands using GDAL. To prevent edge artifacts during inference, we generated continuous 1000 x 1000 pixel image tiles with a 12 pixel buffer across each edge to achieve 1024 x 1024 inputs.

Inputs for the historical imagery tree detection model were more diverse. Channel 1 was grayscale imagery product from the orthorectification step. For channel 2, we calculated gray-level co-occurance matrix (GLCM) variance of the orthoimagery with *ee.Image.glcmTexture* in Google Earth Engine (5-pixel neighborhood, averaging over all four directions). GLCM variance is presented in **equation 1**, and represents the dispersion of pixel value intensity relative to the surrounding neighborhood,

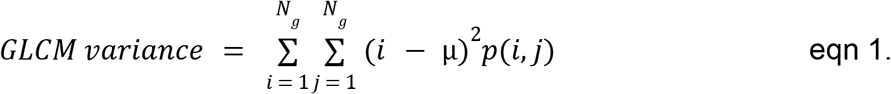

where p(i. j) are coordinates of the spatial-dependence matrix and μ represent the mean of the neighborhood values (Haralick et al., 1973). We rescaled and clamped the GLCM values from [200, 2000] to [0,1] floating point values. Channel 3 represents the dominant land cover classification prior to Euro-american settlement from the Landfire Biophysical Settings datalayer (Landfire BPS v1.4.0). Vegetation cover types were sorted into one of twelve groups based primarily on vegetation structure (e.g., hardwood forest, conifer forest, grassland, shrubland, etc). These classes were then rescaled from [1,12] to [0,1] floating point values. For channel 4, we used the continuous tree cover layer from modern imagery processing, representing the pixel-wise fractional tree cover (%) integrated over one acre. Source data was floating point scaled from [0,1].

#### 2.2.3 Neural Network Training and Inference of Binary Tree Cover Model

We used the Tensorflow 2.5 Python API for training and inference accelerated with an NVIDIA A100 Tensor Core GPU. For training, we used the Dice Coefficient for our loss function (eqn 2). We evaluated 3 standard optimizers provided in Tensorflow (RMSProp, ADAM, and SGD) and achieved the best validation Dice score using the ADAM and RMSProp for the historical and modern tree detection models, respectively. Data augmentation was limited to random rotations of the imagery during training.

We also report the Jacard index (Intersection over Union), sensitivity, and specificity of the model using validation data (**Table 1**).

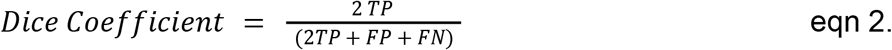

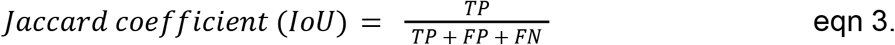

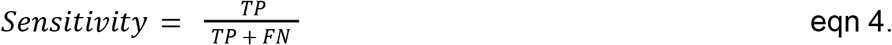

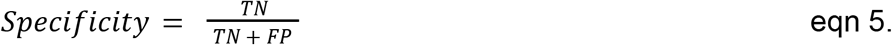

Where TP equals true positive, FP equals false positive, TN equals true negative, and FN equals false negative in a binary classification system. For training and inference, we used a binary threshold value of 0.5.

Inference was performed in Tensorflow 2.5 and the exported image tiles were processed in GDAL 3.4.1 to remove tile buffers and apply affine transformations for reprojection prior to re-ingestion into Google Earth Engine.

#### 2.2.4 Kernel filter and continuous tree cover model

To compare tree cover between historical and modern imagery, we used a kernel filter to convert our binary tree segmentation model outputs into continuous tree cover values. We used the moving window approach developed previously by Falkowski et al. (2017). For each pixel, fractional tree cover was estimated by calculating the mean percent cover over an integrating area of 4046.86 m2 (1-acre). This processing was completed using *ee.Image.reduceNeighborhood(*) with a square kernel in Google Earth Engine.

### 2.3 Mapping areas of tree cover expansion in rangelands

To identify areas of tree cover increase and decrease in rangelands, we first subtracted the historical continuous tree cover model from the modern continuous tree cover model. We then aggregated the difference image to a pixel-size of 30m x 30m to match the scale of the kernel operations. Finally, we applied two filters to identify areas of tree encroachment and tree cover loss. For pixels to score as tree encroachment they needed to have < 2% cover in the historical imagery and > 4% in the modern imagery. To score as tree cover loss, the pixels needed to have > 10% cover in the historical imagery and < 4% in the modern imagery. The 4% threshold was defined operationally, as it was observed to negatively impact tree-sensitive wildlife (Baruch-Mordo et al., 2013). The 10% threshold was chosen as this is the standard tree cover level used by the United States Forest Service (USFS) to define woodlands or forest lands.

#### 2.3.1 Calculating tree expansion area and mapping accuracy

To calculate the area of tree expansion in Montana rangelands, we used the best practice guidelines outlined in (Olofsson et al., 2014) to assess classification accuracy and develop unbiased areal estimates for tree expansion and tree cover loss. We used a random stratified sampling design to evaluate 3,500 pixels based on the relative revalence in the classified map (1000, 500, 2000 30 m X 30 m pixels in areas classified as increasing, decreasing tree cover, or other, respectively). An analyst not involved in the modeling or development of the training data used historical imagery (1 - 2 m GSD), 2019 NAIP imagery (0.6 m GSD), and Google Maps base imagery (0.15 m GSD) in a GIS to evaluate class agreement/disagreement in tree encroachment map. The analysis classified each pixel into one of three classes (tree expansion, tree cover loss, and no change). We developed a count error matrix from the data to calculate bias-corrected areal estimates and confidence intervals. The data presented in this main text represent bias-corrected estimates; performance metrics based solely on the validation data (what is usually reported for ML modeling) are presented in the supplemental.

Following the development of the sampling error (confusion) matrix, we calculated the unbiased estimator 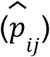 for each cell in the count matrix (**eqn. 6**). Here, *i* refers to the map classification and *j* refers to the reference classification. Each cell classification is weighted by the fractional mapped area for each class (*W_i_*). Here, *n_ij_* represents the count data for a cell in the original error matrix.

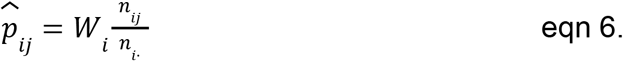

Next, we developed an error-adjusted area estimate for each class 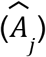 that accounts for omission errors and commission errors over the total map area (*A_tot_*). Here, *q* refers to the number of classes used in the accuracy assessment. This estimate allowed us to estimate the true area of tree expansion after accounting for misclassification in the map product.

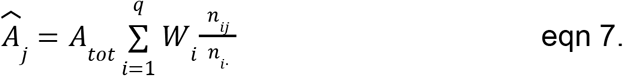

The standard error 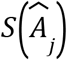 of the error-adjusted estimated area is presented in eqn. 8, with q representing the number of classes in our map. The approximate 95% confidence interval (CI) is calculated by multiplying 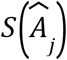 by the z-score (z = 1.96 at 95th percentile).

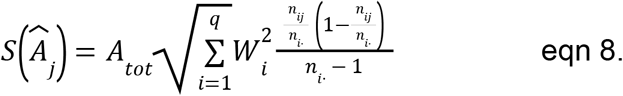

Error-adjusted user accuracy 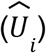 was calculated directly from the estimated error matrix, where 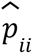 represents the true positive estimator for class i in the map (eqn 9).

This metric is equivalent to precision in standard ML accuracy assessments.

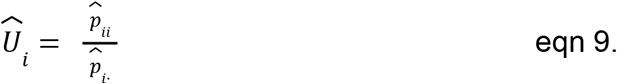

Similarly, error-adjusted producer accuracy 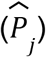 was calculated directly from the estimated-error matrix where 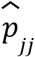 represents the true positive estimator for the reference class. This metric is equivalent to recall in standard ML accuracy assessments.

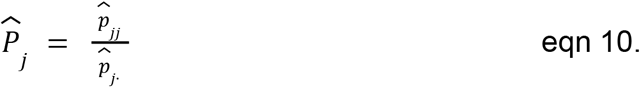

Overall accuracy 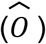 is estimated by summing the true-positive estimators across q classes.

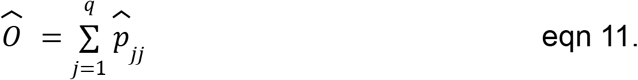

Finally, we conducted a hotspot analysis using the Getis-Ord Gi algorithm to identify regions with high rates of misclassification (Getis & Ord, 1992). We aggregated our point-based misclassification rate into a 0.5 x 0.5 degree grids to identify hotspots at the 95% confidence level.

## 3.0 Results

We processed 17,942 high-resolution historical aerial images for this project. The resulting historical orthoimage product covers 380,832 km^2^; less than 10 km^2^ of imagery was missing from the final product due to missing imagery or processing errors. A total of 1,422 ground control points were placed to assist with georectification. In the final product, we found that the median horizontal (XY) spatial offset was 10 m between historical and modern aerial imagery.

Tree encroachment in Montana grasslands totaled 2,965,587 ± 195,976 hectares between the mid-20th century and 2019, comprising 15.4% of the total modern rangeland area within Montana. The analysis also identified 378,947 ± 86,703 hectares of tree cover loss, covering 2.0% of the rangeland area (**Table 2**). Importantly, these results reflect areal estimates of tree expansion rather than changes in absolute tree cover. Figure 4 shows the extent of tree expansion and tree cover loss across the historical extent rangeland sites in Montana. Map user accuracy (precision) was 88% for tree expansion, indicating that the extent of tree encroachment identified in **Figure 4** is robust. Producer accuracy (recall) was lower (60%), indicating that some tree expansion pixels were missed during mapping. User and producer accuracy for woodland loss were 95% and 32%, respectively. Areas mapped as No Change had similar user accuracy (92%) and producer accuracy (99%). Importantly, the total area estimates and CI for tree encroachment and woodland loss account for producer error, and thus represent unbiased estimates of land cover change.

**Table 2:** Tree encroachment in Montana grasslands determined from analysis of historical and modern aerial imagery.

|  | **No change detected** | **Tree cover expansion** | **Tree cover loss** |
| --- | --- | --- | --- |
| Area (ha) | 15,927,228 | 2,965,587 | 378,947 |
| 95% CI (ha) | 210,722 | 195,976 | 86,703 |
| Standard error (ha) | 107,511 | 99,988 | 44,236 |
| User accuracy (precision) | 0.92 | 0.88 | 0.95 |
| Producer accuracy (recall) | 0.99 | 0.60 | 0.32 |
| F1-Score | 0.95 | 0.71 | 0.47 |

**Figure 4:**
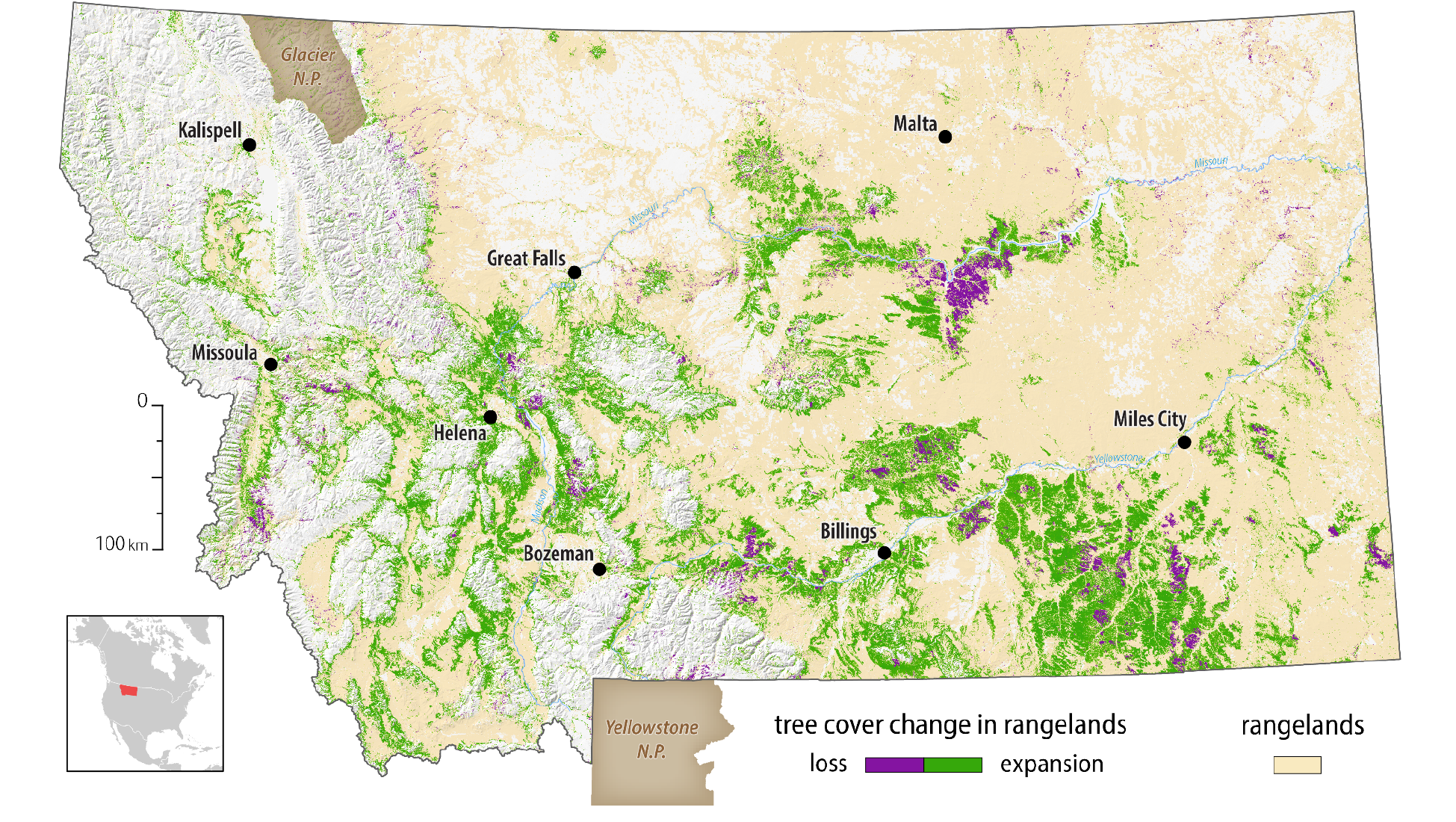
Tree cover expansion and loss across Montana rangelands. Forests, agricultural lands, wetlands, and the built environment were excluded from the analysis.

We summarize tree expansion and tree cover loss by ecosystem type in **Table 3**. Grasslands were most impacted by acreage (1,416,665 ± 93,618 ha, 11.3% of area), but we saw greater impacts to shrublands and woodlands (20.7% and 33.1% tree expansion by area, respectively). Tree cover loss was found to be occurring across 1.0% of grasslands and shrublands, but losses were much higher in open woodlands (16.3% of area).

**Table 3:**
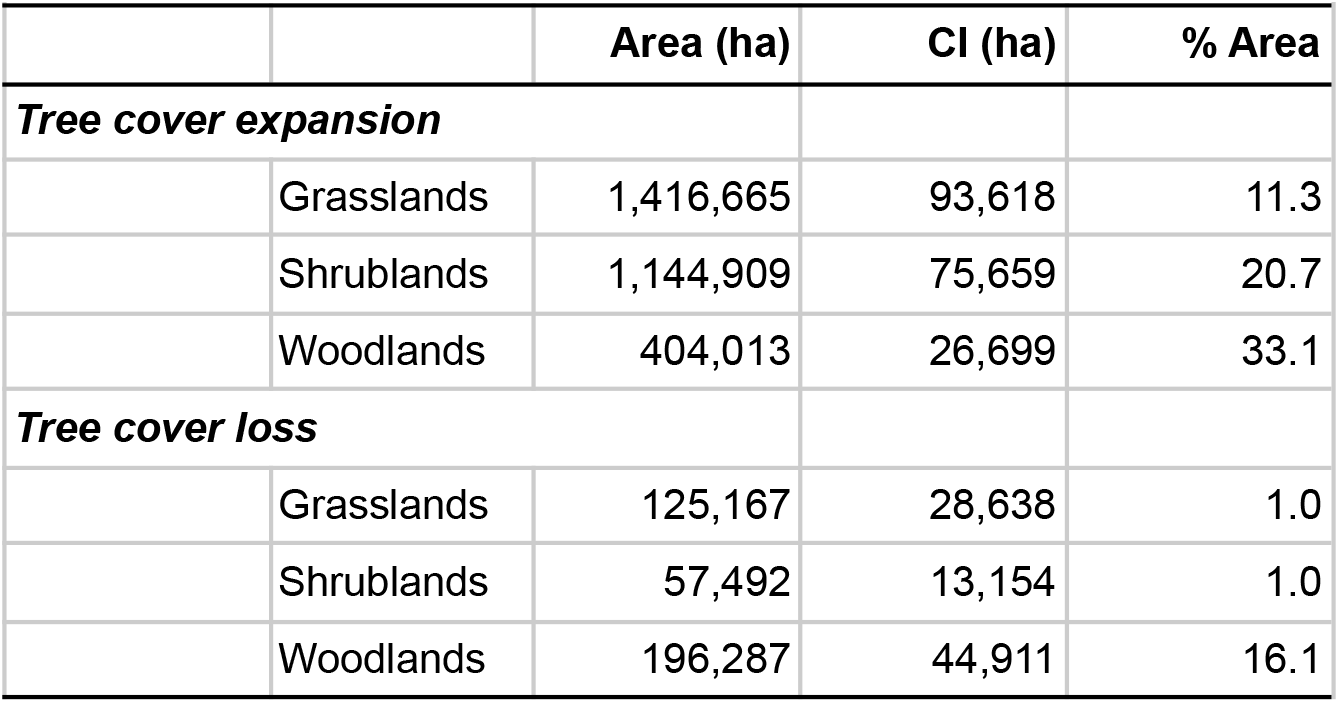
Tree cover change metrics by ecosystem type

To identify areas of tree expansion and woodland loss missed by our model, we performed a hot-spot analysis to identify and investigate areas where tree encroachment and woodland loss were misclassified. **Figure 5** shows the spatial distribution of points used in our accuracy assessment and misclassification hotspots based on the Getis-Ord Gi algorithm. We identified seven possible hotspots at the 95% confidence level. Based on visual examination, two of the seven areas had classification errors that would misrepresent land cover change to map users. These areas are labeled in **Figure 5** and are located in the eastern half of the Northern Cheyenne Indian Nation. Misclassification was attributable to the failure of the U-Net model to identify tree cover in the historical imagery due to low-contrast source imagery.

**Figure 5:**
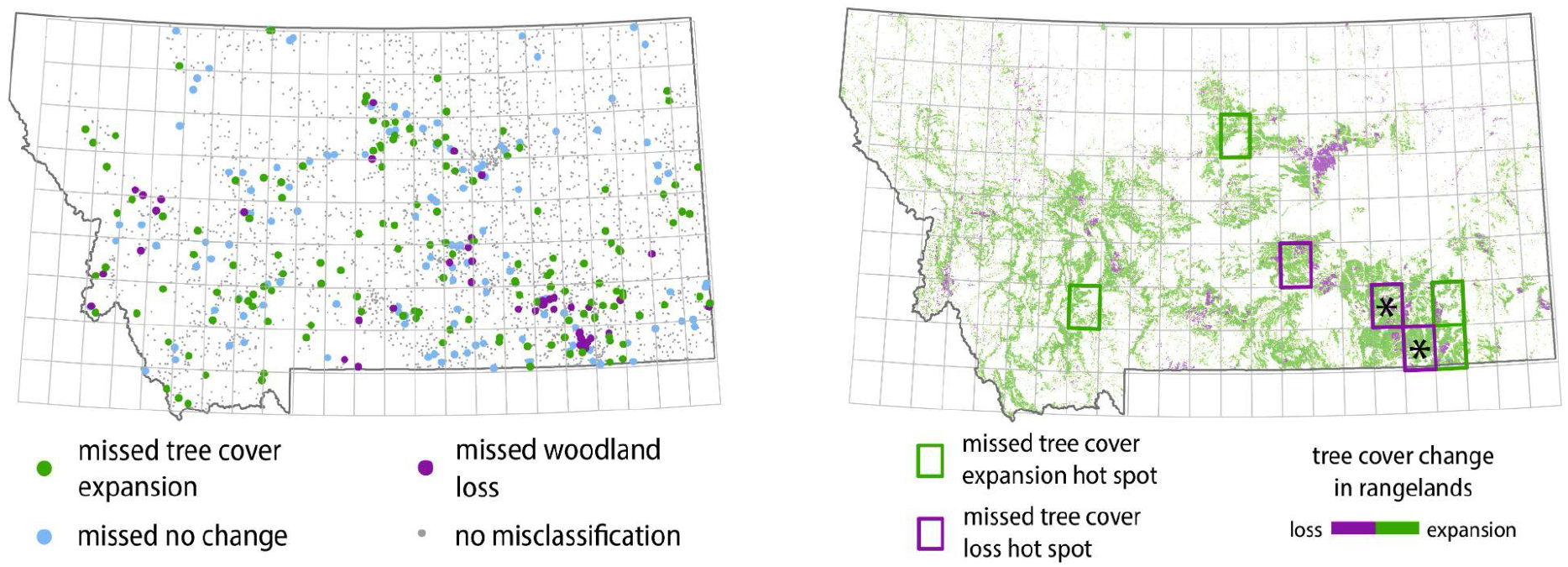
Accuracy assessment. Left panel shows 3,500 points analyzed for model accuracy. Right panel shows areas identified as misclassification hotspots using the *Getis ord Gi* algorithm; Post hoc analysis of hotspots found two areas of gross misclassification due to localized failure of the tree-detection algorithm (identified with asterisk).

The tree feature detection model used in this analysis was successful in segmenting trees in rangelands, woodlands, and most forested ecosystems in both modern and historical imagery. For the historical imagery model, sensitivity and specificity in our validation dataset were found to be 0.9858 and 0.9855, respectively. We found that the U-Net architecture appeared to have had difficulty with false positive classifications in agricultural areas, more mesic (wet) sites, and areas of higher textural complexity (e.g., rock outcrops with shadows) in the historical imagery (Fig. 6). However, these areas of misclassification generally had little impact on the overall analysis; agricultural areas and wetlands were masked in the final analysis, and mesic sites in uplands generally did not result in tree cover estimates greater than 10%.

**Figure 6:**
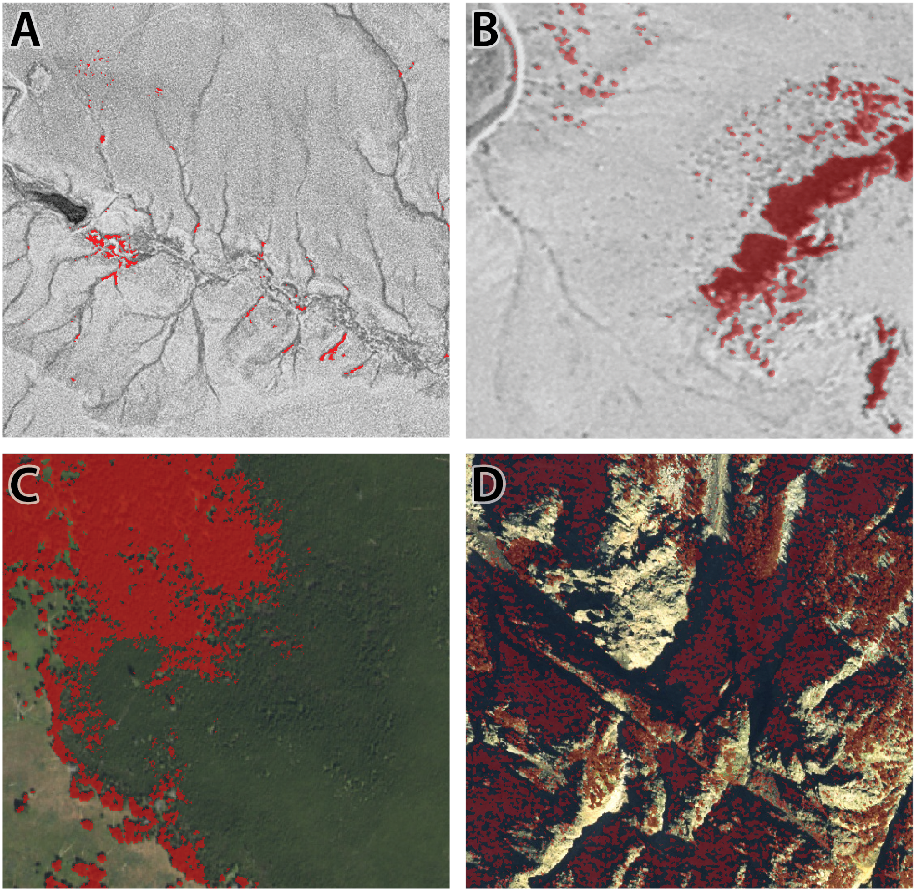
Examples of common feature detection errors where red areas are misclassified as trees. Treeless wet drainage areas and rock outcrops were commonly classified as trees in historical imagery (panels A and B). In modern imagery, thick lodgepole pine forests were often misclassified (Panel C), as well as shadowed areas in high-relief alpine locations (Panel D).

The U-Net model was also successful at segmenting tree features in modern NAIP imagery (sensitivity: 0.9314, specificity: 0.9769), except for areas of complete and uniform tree cover (e.g. *Pinus contorta* stands) where false negative detections were common (Fig. 6). However, because primary forests were omitted from our analysis of tree expansion in rangelands, this misclassification had little impact on identifying tree expansion among rangelands. Areas in deep shadow and on high slope were also misclassified, but these areas were less common among mapped rangelands.

## 4.0 Discussion

We found that more than 15% of Montana’s rangelands have experienced tree expansion since the mid-20th century, with the majority of expansion occurring among grassland and shrubland sites (2.56 million ha). These findings are consistent with other analyses of tree expansion in the western United States and the recent acceleration of tree expansion in the northern Great Plains (Filippelli et al., 2020; Morford et al., 2022a). Roughly 33% of woodlands were experiencing tree expansion, but these sites also saw relatively higher tree cover loss (16.1% of area), suggesting that disturbance mechanisms (fire, drought, and beetle kill) were more effective at regulating tree expansion among these sites vs. grasslands and shrublands. Considering expansion and loss processes together, Montana’s shrubland ecosystems appear most impacted by tree expansion, aligning with broader, biome-wide analyses showing rapid declines in the extent of intact sagebrush habitats across the western United States (Doherty et al., 2022).

Compared to satellite-based analyses, a baseline from the mid-20th century captures an additional four decades of ecosystem change. Using this earlier baseline, we quantified roughly twice as much tree expansion in Montana grasslands and shrublands as can be quantified using more recent satellite data (Fig. 7, Morford et al., 2022). In some cases, we discovered that our historical imagery modeling approach is more sensitive to incipient tree expansion. Despite these differences, we found similar broad spatial patterns of tree expansion using aerial and satellite imagery, implying that satellite-based methods are similarly effective at identifying where tree expansion is occurring on the landscape—albeit with a later baseline.

**Figure 7:**
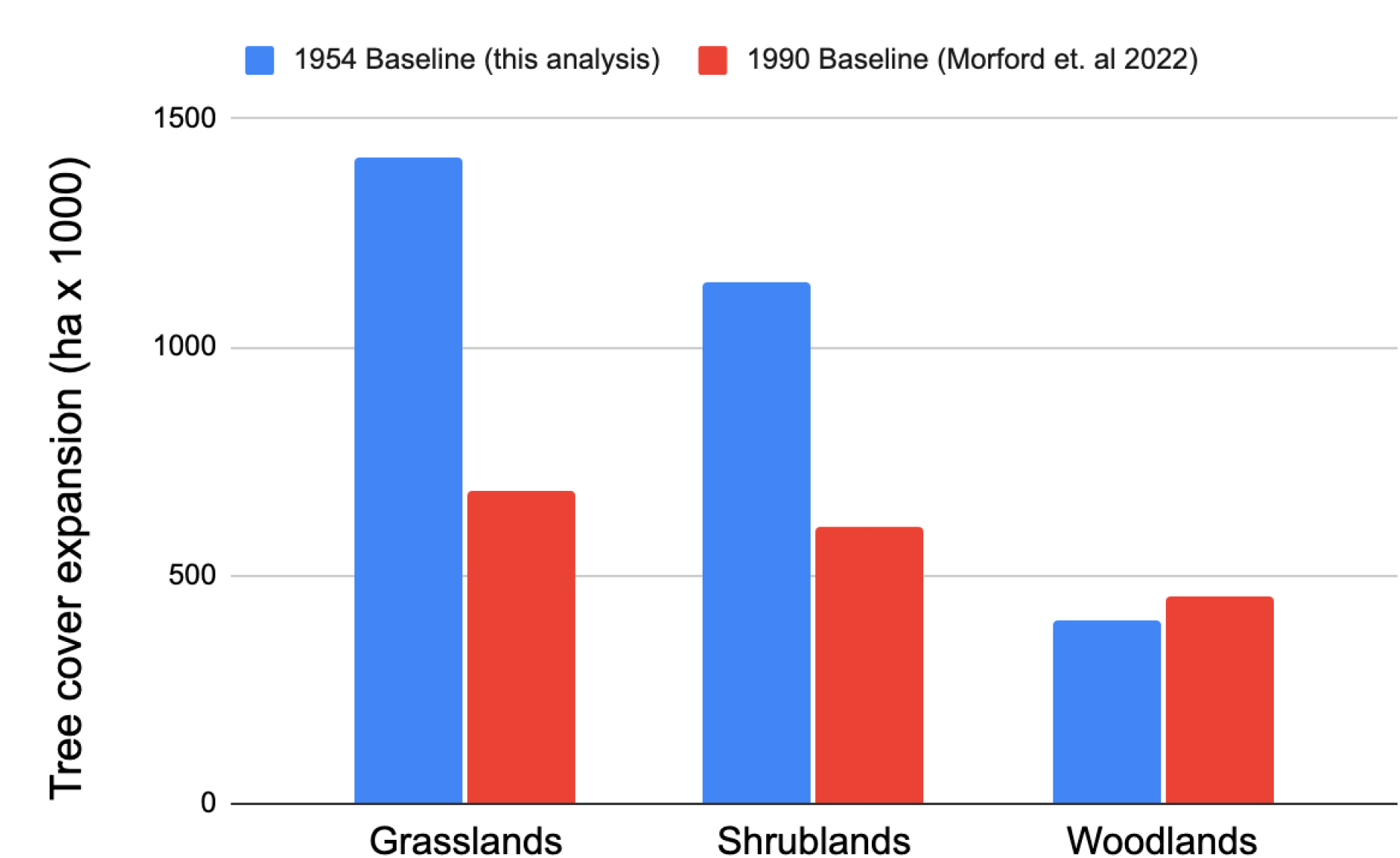
Comparison of tree cover expansion estimates using historical imagery (this analysis) and satellite imagery (Morford et. al 2022). The historical imagery approach identified nearly double the amount of tree cover expansion in grasslands and shrublands. These differences are attributable to differences in baselines (1954 vs. 1990) and improved detection of tree recruitment using aerial imagery.

Wildfire was responsible for most of the tree cover loss in Montana’s rangelands. We intersected our tree cover loss map with fire perimeter data from 1984 - 2019 and found that 81.3% of tree cover loss corresponded to lands burned in large wildfires (Eidenshink et al., 2007). In burned areas, we often saw standing dead trees among grasslands and shrublands recovering from the fire. Even when fire successfully eliminated seed sources and prevented the growth of new trees, the existing tree structures that contribute significantly to the loss of animal biodiversity (i.e., standing dead trees that provide hunting perches for predators) remained. This suggests that increasing fire frequency in the western United States may not immediately improve biodiversity outcomes in rangeland ecosystems where tree expansion has occurred, even if other ecosystem functions and biogeochemical cycling recover as a result of fire disturbance (Burke et al., 2021; Williams et al., 2020).

Mapping landscape-scale tree expansion using deep learning models and historical aerial imagery was an effective strategy for quantifying change over time periods that are unavailable with satellite data. However, because of variability in data quality and inconsistent imagery orthorectification, this approach introduces additional modeling challenges that do not exist when using curated satellite imagery, such as Landsat Collection 2 (Micijevic et al., 2020). Differences in contrast and vignetting in the historical imagery cause issues with tree feature detection during inference that are difficult to detect and mitigate using our semi-automated approach. Furthermore, XY offset in historical imagery orthorectification contributes to biased detection of tree expansion and tree cover loss in some cases, particularly near woodland edges (**Fig. 8**). These issues, however, can likely be addressed with improved imagery restoration and deep-learning assisted georectification (Feng et al., 2021; Su et al., 2022).

**Figure 8:**
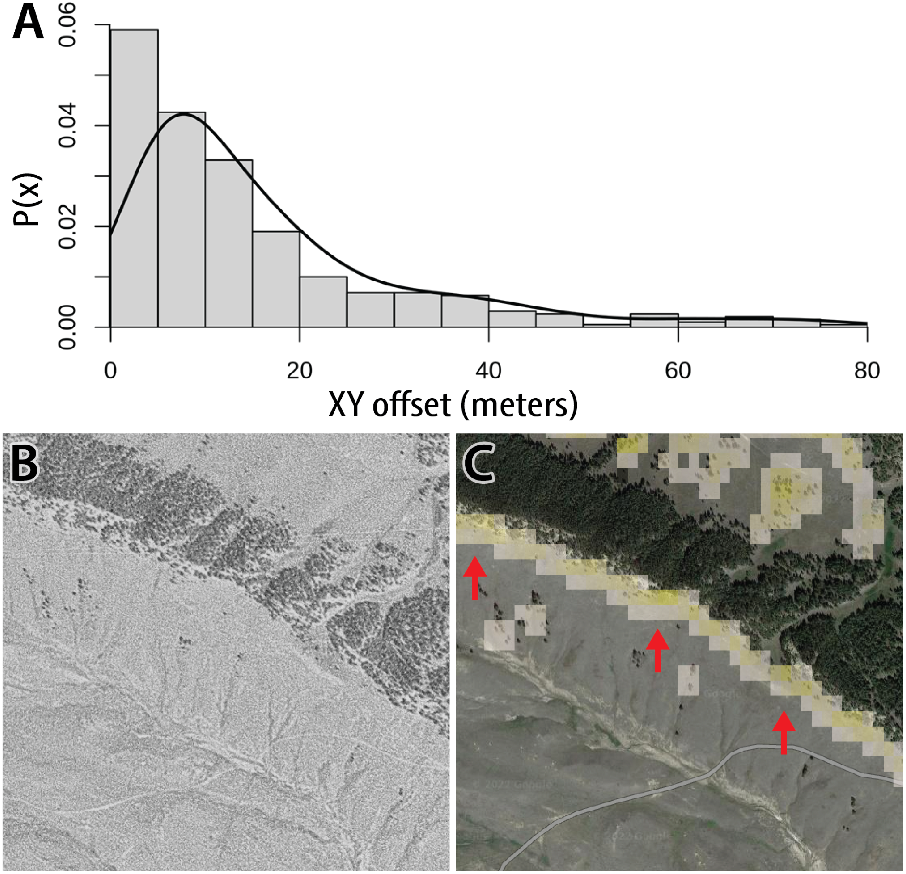
Horizontal (XY) offset between the historical imagery and modern imagery was generally 10 m or less (A). Panels B and C show how high offset can result in the misclassification of tree cover expansion (in yellow along linear features. Arrows point to misclassification along the forest edge.

Our findings have implications for conservation-focused tree encroachment management in U.S. grasslands and shrublands. Given the scope and extent of this environmental change, our results suggest that management strategies that fail to consider the landscape-level trajectory for tree encroachment will likely be ineffective at achieving durable grassland conservation. Small-scale, isolated, tree removal projects in highly encroached areas contribute little to grassland conservation if trees and tree seed sources persist nearby (Twidwell et al, 2021). Instead, conservation managers should prioritize addressing early tree expansion in high-quality, but vulnerable, intact grasslands and shrublands (Maestas et al., 2022; Morford et al., 2022; Yokomizo et al., 2009). For example, combining our tree expansion maps with conservation priority area maps can help prioritize where conservation funds should be invested to have the greatest likelihood of achieving desired outcomes at a landscape scale (**Fig. 9**).

**Figure 9:**
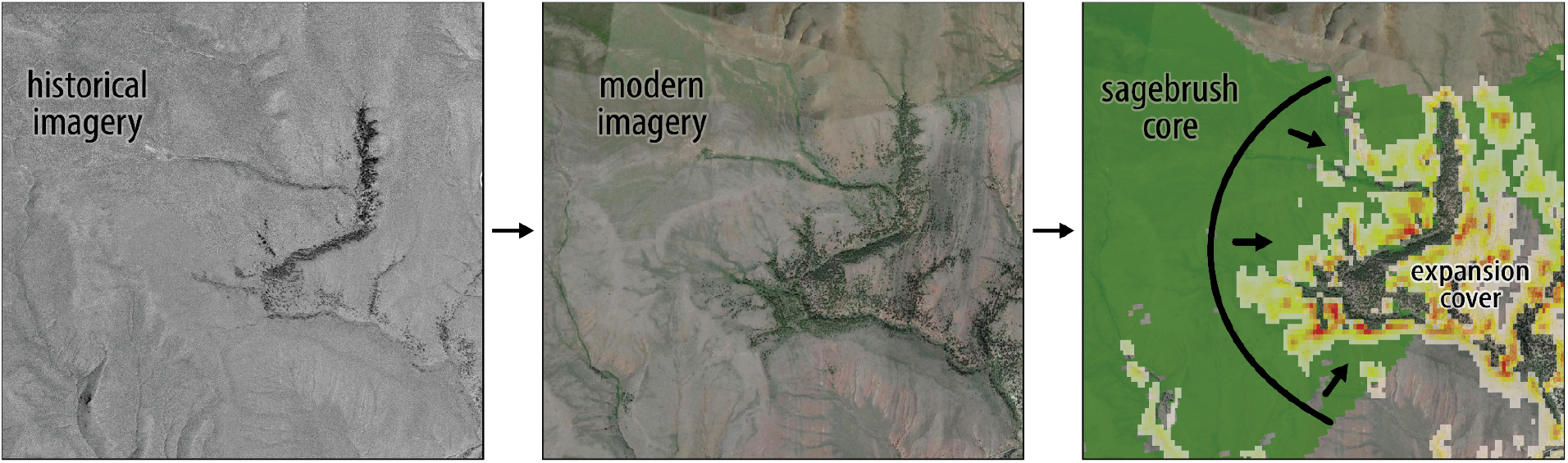
Conservation strategies should leverage historical imagery with an understanding of where core-intact grassland and shrubland biomes intersect areas of tree expansion. Conserving intact biomes should be prioritized, with management activities focused first on removing invading trees and seed sources from intact core areas rather than removal of trees from highly invaded areas. (sagebrush core areas from Doherty et al., 2022).

The tree expansion map and historical imagery provide a compelling conservation tool for communicating how tree expansion reorganizes the structure of rangeland ecosystems. The slow pace of tree expansion and its impacts to ecosystem function and biodiversity are difficult to observe directly, especially in the early stages of tree colonization when conservation management is most effective. Presenting a visual landscape-level view of tree expansion in an easy-to-use mapping decision support tool can help overcome some of the limitations of shifting baseline syndrome (Soga & Gaston, 2018). Notably, historical imagery does not represent reference conditions at every location, so caution should be exercised when attempting to infer localized process-level change. We propose combining historical imagery with land use and disturbance history data to make informed management decisions that integrate landscape-level tree expansion trajectories with local-scale site conditions and conservation priorities. We provide a web application to visualize historical imagery and tree expansion in Montana rangelands to facilitate far-reaching use of these data.

## 5.0 Conclusions

Tree expansion is rapidly altering the structure of grasslands in Montana and around the world. Converting grasslands and shrublands to forests reduces the extent of these most threatened biomes, causing biodiversity loss to accelerate. This imagery analysis and mapping product provide an intuitive way to visually and numerically convey the extent of tree encroachment and its effects on ecosystem structure. Management practitioners can use our map and supporting historical imagery tool to support development of targeted conservation plans to address this environmental challenge.

## 6.0 Acknowledgements

This work was made possible by the USDA-Natural Resources Conservation Service’s (NRCS) Conservation Collaboration Grant (Agreement NR200325XXXXC002). NVIDIA provided hardware used in this analysis through their Academic Hardware Grants program. The findings and conclusions in the publication are those of the authors and should not be construed to represent the views of the USDA, U.S. Fish and Wildlife Service, or the U.S. Government. Any use of trade, firm, or product names is for descriptive purposes only and does not imply endorsement by the U.S. Government. The authors declare no conflicts of interest.

---

**Supplemental Table 1:**
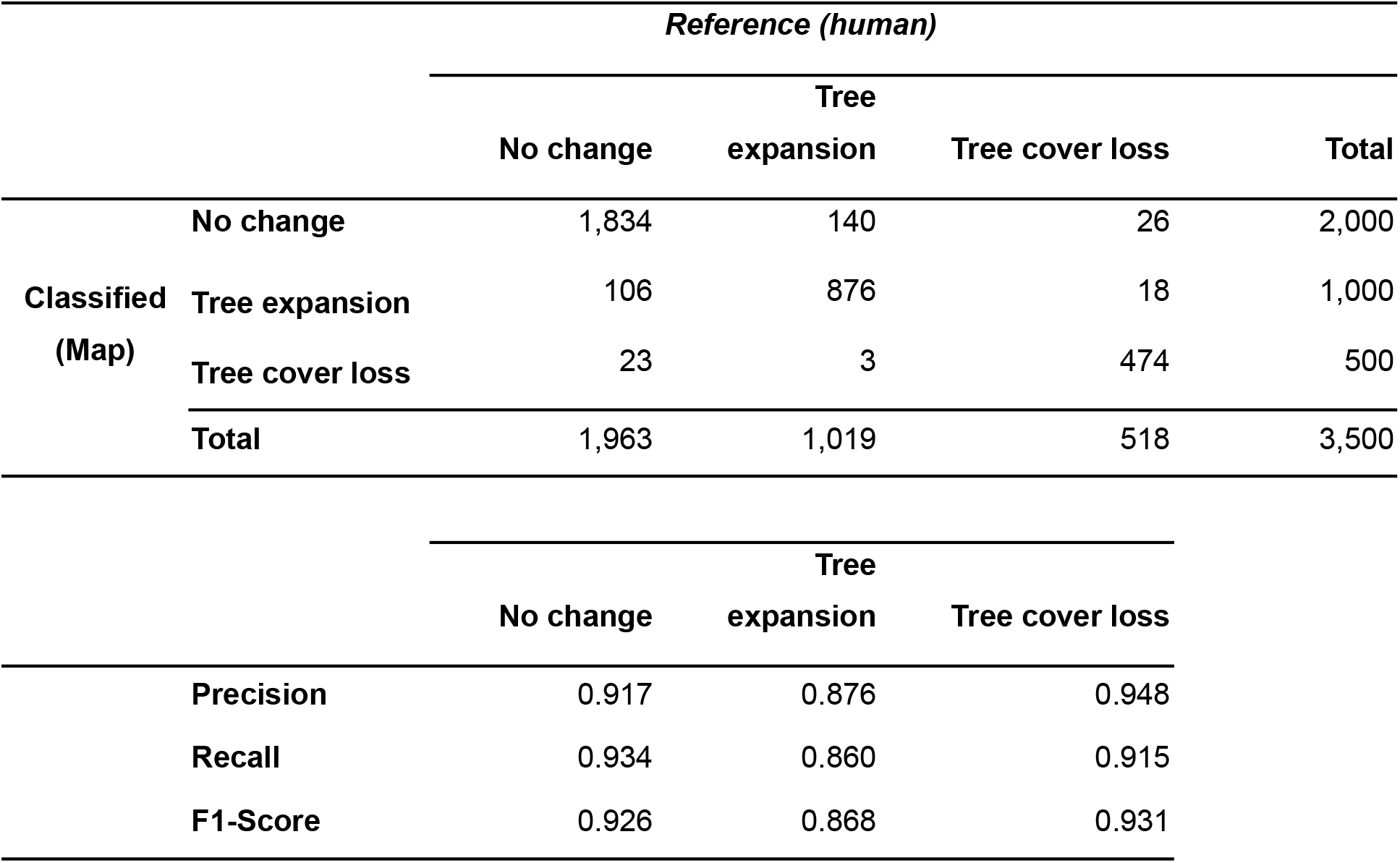
Confusion Matrix and classification metrics from raw accuracy analysis.

